# Nascent RNA kinetics with complex promoter architecture: Analytic results and parameter inference

**DOI:** 10.1101/2023.12.28.573588

**Authors:** Changhong Shi, Xiyan Yang, Tianshou Zhou, Jiajun Zhang

## Abstract

Transcription is a stochastic process that involves several downstream operations, which make it difficult to model and infer transcription kinetics from mature RNA numbers in individual cell. However, recent advances in single-cell technologies have enabled a more precise measurement of the fluctuations of nascent RNA that closely reflect transcription kinetics. In this paper, we introduce a general stochastic model to mimic nascent RNA kinetics with complex promoter architecture. We derive the exact distribution and moments of nascent RNA using queuing theory techniques, which provide valuable insights into the effect of the molecular memory created by the multistep activation and deactivation on the stochastic kinetics of nascent RNA. Moreover, based on the analytical results, we develop a statistical method to infer the promoter memory from stationary nascent RNA distributions. Data analysis of synthetic data and a realistic example, the *HIV-1* gene, verifies the validity of this inference method.

## Introduction

Transcription is a complex biochemical process that leads to stochastic fluctuations in mRNA and subsequent protein production. The mature mRNA distribution data can be modeled using a two-state gene expression model, which has successfully revealed the mechanism of bursting and the non-Poisson nature of transcription [1-7]. However, the expression levels of mature mRNA are strongly influenced by various post-transcriptional processes such as splicing, nuclear export, and miRNA mediation [8-10]. Therefore, conventional gene-expression models focusing on mature mRNA may not be suitable for interpreting and inferring transcriptional processes, and more elaborate models need to be developed.

An alternative to measuring transcriptional products is to count the number of nascent RNAs (or the number of RNA polymerase molecules, RNAP) at the single-cell level. Recent experimental methods such as molecular fluorescence in situ hybridization (smFISH) can provide a steady-state distribution of nascent RNA [11-13]. Fitting experimental data of nascent RNA using stochastic models of nascent RNA allows for a more precise understanding of the regulatory steps involved in transcriptional processes [14, 15]. Therefore, numerous stochastic models have been recently developed to simulate and infer nascent RNA kinetics [16-23]. For instance, Choubey, et al., [22] proposed a two-state model for the stochastic kinetics of nascent RNA, which assumed that a gene switches between an active and an inactive state, transcription initiation occurs only in the active state, and polymerase moves from one base pair to the next until reaching the end of the gene. The authors used queuing theory to derive the analytical expressions of transient and stationary expectations, as well as the variance of nascent RNA in the case of deterministic elongation time. Additionally, Heng, et al., [14] introduced a coarse-grained version of that model by neglecting RNAP movement between base pairs and derived the analytical expression of steady-state nascent RNA distribution.

It is worth noting that the two-state model assumes that the time spent in the active or inactive state follows an exponential distribution. While this assumption may be reasonable for bacteria [24], recent experimental studies have shown that the inactive periods of many genes in mammalian cells follow non-exponential distributions [25]. This suggests that the transition from an inactive to an active state is a multi-step process containing multiple inactive states. More generally, multiple transcriptional active and inactive states may exist in both prokaryotic and eukaryotic cells. For example, in *E. coli*, the Promoter for Repressor Maintenance (PRM) promoter of phage lambda is regulated by two different transcription factors (TFs) binding to two sets of operators, which can be brought together through DNA looping. Consequently, the number of regulatory states of the PRM promoter is up to 128 [26]. In contrast, eukaryotic promoters would be more complex since they involve nucleosomes competing with or being removed by TFs [27]. In addition to the conventional regulation by TFs, eukaryotic promoters can undergo epigenetic regulation through histone modifications [28-30], leading to complex promoter structures. A multi-step process can create the memory between molecular events, implying that waiting-time distributions between these events are non-exponential and lead to non-Markovian dynamics of the underlying system.

In recent years, gene expression models with complex promoter multi-state switching mechanisms have been widely studied theoretically [31-41]. However, most of these studies focus on the distributions of mature RNA and protein in a steady-state or transient state, and how promoter structure affects nascent RNA kinetics remains incompletely understood. Recently, several stochastic models of nascent RNA with multi-state promoter structure have been studied. Holehouse et al. [19,57] proposed a mechanistic model with 3-state promoter structure and obtained exact steady state distribution for nascent RNA using the delayed chemical master equation (dCME). Juraj et al. [18] developed a general framework that allowed the derivation of nascent RNA distributions for gene models with stochastic and multistep initiation but with deterministic elongation and termination. This framework is based on the assumption that the transcription initiation is a renewal process, which is true when there is only one active gene state and multiple inactive gene states. Also, we note that previous studies primarily examined the steady-state or time-dependent behavior of mature mRNA or protein. Thus far, there has been still a lack of detailed studies on stochastic nascent RNA kinetics in the case of arbitrarily complex promoter structures. Therefore, the aim of this study is twofold: to develop a general stochastic model of nascent RNA with complex promoter structure and to derive the nascent RNA distribution to show how molecular memory impacts the stochastic kinetics of nascent RNA.

### A general stochastic model of nascent RNA kinetics

We introduce a stochastic gene-expression model with complex promoter switching followed by a deterministic elongation process. Assume that the gene promoter has *L* distinct states, denoted by *S*_1_, *S*_2_,…, *S*_*L*_ . Biophysically, these states correspond to different conformational states during chromatin remodeling or to different binding states for TFs. We use the matrix **W** = [*ω*_*ij*_] to describe the switching rates between promoter states, where elements *ω*_*ij*_ represent the transition rate from state *S*_*i*_ to state *S* _*j*_, satisfying the relationship 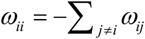. We also use the diagonal matrix **Λ** = diag(*k*_1_, *k*_2_,…, *k*_*L*_) to describe the rates of transcription initiation, where *k*_*i*_ is the transcription initiation rate when the promoter state is at *S*_*i*_ state. Also assume that the elongation of a single transcript is deterministic and the elongation speed is a constant *k*_elo_ kb/min. Consequently, the elongation time is *t*_elo_ = *l*_gene_ *k*_elo_, where *l*_gene_ is the gene length. Although a transcriptional elongation process would be very complex, e.g., it would involve pausing and backtracking of polymerases along the gene body, our assumption of deterministic elongation is justified for a vast number of genes [22,42]. We set the survival function of the nascent RNA as

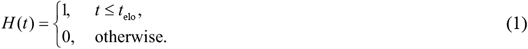

Let *P*_*i*_ (*n,t*) denote the probability that the number of nascent RNAs is *n* at time *t*, given that promoter state is *i* at time 0 and the number of nascent RNA is 0 at time 0, i.e.,

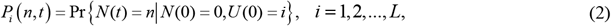

where *U* (*t*) and *N* (*t*) represent the promoter state and the nascent RNA number at time *t*, respectively. For convenience, we denote ***P*** (*n,t*) = (*P*_1_ (*n,t*),…, *P*_*L*_ (*n,t*))^T^, where the symbol ‘T’ represents transpose. Using the method proposed in our recent study [43], we can derive the following so-called generalized master equation for ***P***(*n,t*)

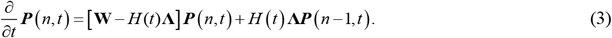

Let ***G*** (*z,t*) = (*G*_1_ (*z,t*),*G*_2_ (*z,t*),…,*G*_L_ (*z,t*))^T^ be the probability generating function corresponding to ***P*** (*n,t*), i.e., 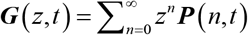. Then Eq. (3) can be transformed to

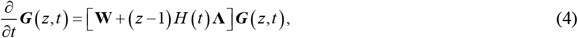

with the initial condition ***G*** (*z*,0) = ***e***, where ***e*** = (1,1,…,1)^T^.

As a special case, we consider the model with a cyclic promoter structure, i.e., the promoter proceeds sequentially through several irreversible active (ON) state and inactive (OFF) states, which altogether form a cycle (Fig. 1 (A)). Note that this cyclic model can be used to describe many processes involved in transcription such as recruitment of various polymerases and transcription factors, assembly of the pre-initiation complexes, chromatin remodeling, and histone modification [36]. For simplicity, we assume that the transcription initiation rate is *k*_1_ at each active state and *k*_0_ at each inactive state, where *k*_1_ > *k*_0_ . Specifically, we assume the gene promoter has *L* states, which are distinguished into *L*_1_ active states and *L*_2_ inactive states. Denote by 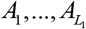 ON states and by 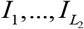 OFF states. The corresponding biochemical reactions are listed in Table 1. Note that if *L*_1_ = 1 and *L*_2_ > 1, the cyclic model becomes the multi-OFF mechanism (i.e., the promoter has multiple OFF states but only has one ON state). If *L*_1_ > 1 and *L*_2_ = 1, the cyclic model becomes the multi-ON mechanism (i.e., the promoter has multiple ON states but only has one OFF state). In particular, if *L*_1_ = 1 and *L*_2_ = 1, the cyclic model reduces to the classical two-state model. In a word, the cyclic model includes many transcription models being previously studied.

**Table 1.** Biochemical reactions for a transcription model to be studied.

| Reaction | Description |
| --- | --- |
| $A_i \xrightarrow{\omega_{i,i+1}^{(on)}} A_{i+1}, i = 1, 2, \dots, L_1 - 1$ | Transitions between active states |
| $I_j \xrightarrow{\omega_{j,j+1}^{(off)}} I_{j+1}, j = 1, 2, \dots, L_2 - 1$ | Transitions between inactive states |
| $A_{L_1} \xrightarrow{\omega_{L_1,1}^{(on)}} I_1, I_{L_2} \xrightarrow{\omega_{L_2,1}^{(off)}} A_1$ | Transitions between active and inactive states |
| $A_i \xrightarrow{k_1} A_i + \text{nascentRNA}$ | Transcription initiation at the active state |
| $I_j \xrightarrow{k_0} I_j + \text{nascentRNA}$ | Transcription initiation at the inactive state |
| $\text{nascent RNA} \xrightarrow[t_{\text{elo}}]{} \emptyset$ | Nascent RNA elongation |

**Fig. 1.**
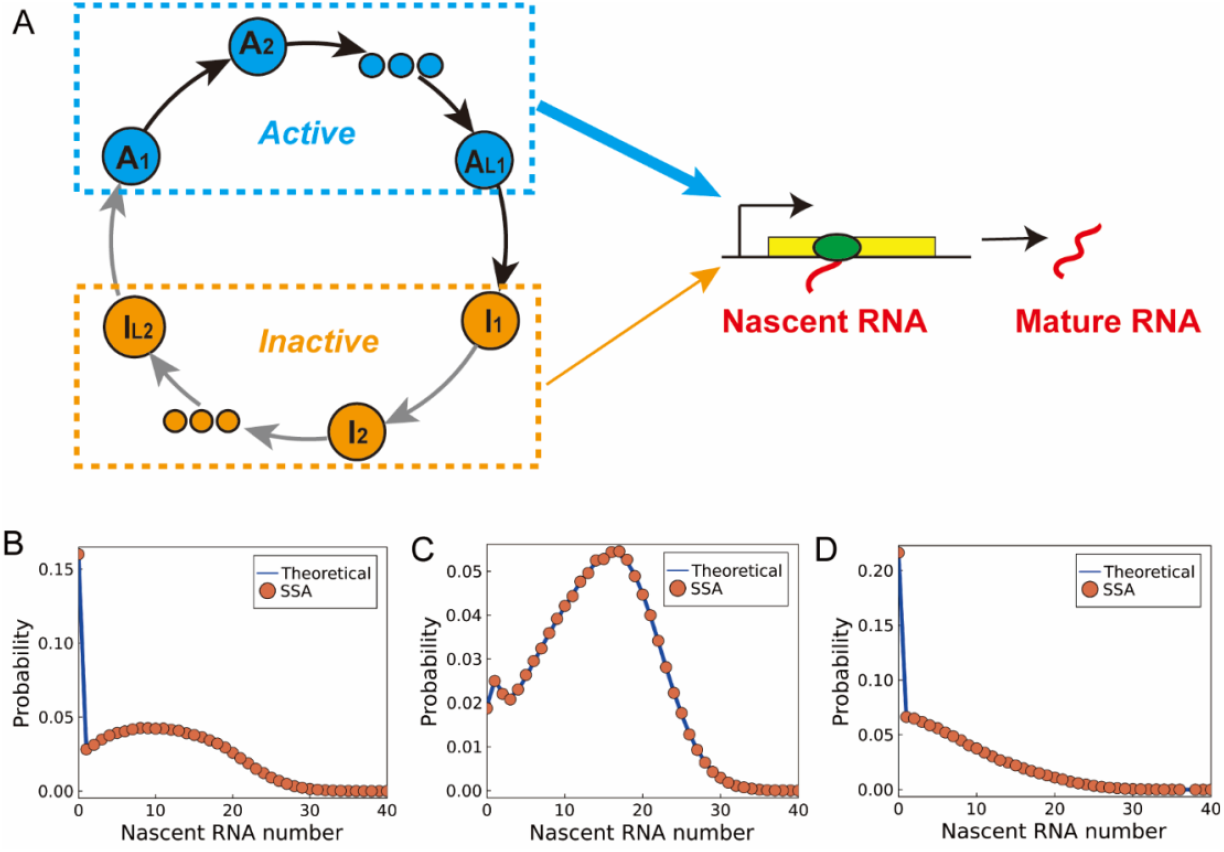
Schematic diagram for a model of nascent RNA with the cyclic promoter structure, where the blue and red cycles represent transcription active (ON) and inactive (OFF) states, respectively. (B)(C)(D) The steady-state nascent RNA distributions obtained by the matrix-form solution and the stochastic simulation algorithm (SSA). The parameter values are set as: (A) *u* = 1,*v* = 1.5, *L*_1_ = *L*_2_ = 2, *k*_1_ = 20, *k*_0_ = 0. (B) *u* = 0.7, *v* = 1.5, *L*_1_ = *L*_2_ = 2, *k*_1_ = 20, *k*_0_ = 1. (C) *u* = 2,*v* = 1.5, *L*_1_ = 1, *L*_2_ = 3, *k*_1_ = 20, *k*_0_ = 0.

To make an equitable comparison between models with different values of *L*, we assume that the transition rates at the ON (OFF) states are the same, i.e., 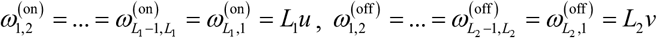, where *u* and *ν* are two constants. With these assumptions, the dwell time at the ON state, *τ* _on_, obeys an Erlang distribution with shape parameter *L*_1_ and mean 1 *u*, and the dwell time at the OFF state, *τ*_off_, also follows an Erlang distribution with shape parameter *L*_2_ and mean 1 *ν* . Note that if the noise of the dwell time is characterized by the coefficient of variation, it is 1 *L*_1_ for the dwell time *τ* _on_, and 1/*L*_2_ for the dwell time *τ*_off_ . As *L*_*i*_ → ∞, this noise tends to zero and the dwell time converges to a deterministic value and the molecular memory is the strongest in each case. In the following, *L*_1_ and *L*_2_ are called as memory index of promoter activation and deactivation, respectively.

### Distribution of nascent RNA in steady state

In a previous study [43], we used the finite state projection (FSP) method to calculate a numerical nascent RNA distribution ***P***(*n,t*) based on Eq. (3). Although this method can ensure that the approximation error is within a range of some pre-specified tolerance level, it is inefficient to obtain an accurate nascent RNA distribution in the steady state since it often takes a long time when a system reaches the steady state. In this section, we present two forms of the exact solutions for the stationary distribution of nascent RNA: Laplace-form solution and matrix-form solution.

### Laplace-form solution

The closed-form solution of Eq. (4) is generally difficult to obtain. Under the assumption that the elongation time is deterministic, we find that the solution of Eq. (4) takes the form

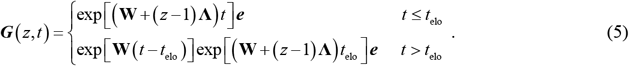

Note that matrix exponential function exp[**W**(*t* ‡ *t*)] tend to ***eπ*** when *t* → ∞, where the row vector ***π*** denotes the steady state of the promoter, satisfying ***π*W** = **0** and ***πe*** = 1 . Since the steady-state distribution of nascent RNA is independent of the initial state: *P* (*n*) ≡ *P*_*1*_ (*n*, ∞) = … = *P*_*L*_ (*n*, ∞), we have *G*_1_ (*z*) ≡ *G*_2_ (*z*, ∞) = *G* (*z*, ∞) = … = *G*_*L*_ (*z*, ∞) .

Combining these results with Eq. (5), we can derive the analytical expression of the stationary nascent RNA distribution as follows

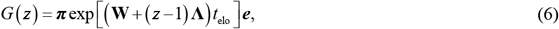

which is a function of *t*_elo_ in principle, re-denoted by *G* (*z,t*_elo_) . The Laplace transform of *G* (*z,t*_elo_) with respect to *t*_elo_ is given by

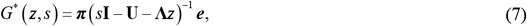

where **U** = **W** ‡ **Λ** . Note that the function *G*^*^ (*z, s*) has the following Taylor expansion

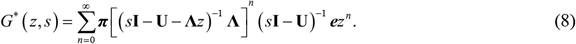

On the other hand, according to the definition of probability generating function, we know that function *G*^*^ (*z, s*) can be written as 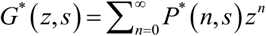 where *P*^*^ (*n, s*) is the Laplace transform of *P*(*n*) (which is also a function of *t*_elo_ in principle) with respect to *t*_elo_ . Therefore, we obtain the Laplace-form solution of stationary nascent RNA distribution

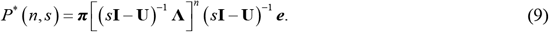

Then, the inverse Laplace transform gives the explicit expression of the probability distribution have

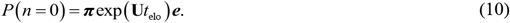

For *n* > 0, it is possible, albeit tedious, to derive the analytical expression of *P*(*n*) . In any case, we can numerically calculate the probability distribution with the above formula.

We can also derive the analytical expression for the stationary moments of the nascent RNA with a general promoter switching. Let *u*^*(k)*^ denotes the *k-*th factorial moments of nascent RNA number *N* in the steady state, i.e., *u*^*(k)*^ = *E* [*N* (*N* ‡1)…(*N* ‡ *k* +1)] . Applying the formula 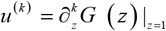 to Eq. (6), we obtain the first two factorial moments (see refs. [44,45] for details)

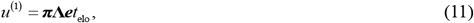

and

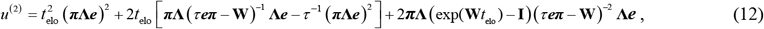

where *τ* ≠ 0 . Therefore, the Fano factor, which is defined as the ratio of the variance over the mean, is given by

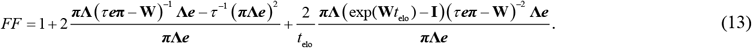

Note that the value of *FF* does not depend on the choice of *τ* . Setting *τ* = ‡1, we have

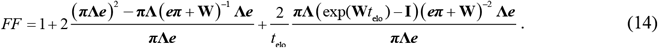

In particular, as *t*_elo_ → ∞, we have

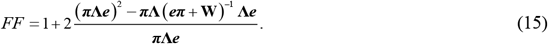

### Matrix-form solution

Although Eq. (9) allows us to calculate the closed-form solution of stationary nascent RNA distribution for some special cases, we find that it is inefficient to calculate the stationary distribution numerically in general cases. Therefore, we try to derive the matrix-form solution of the stationary nascent RNA distribution to facilitate numerical computation. Assuming that there exists a large *N* such that *P*_*i*_ (*n,t*) = 0 for *i* = 1, 2,…, *K* and *n* > *N* +1. We denote

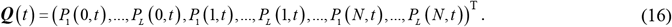

For *t* ≤ *t*_elo_, Eq. (3) can be solved by the following ordinary differential equation (ODE) up to time *t*

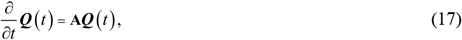

where matrix **A** takes the form

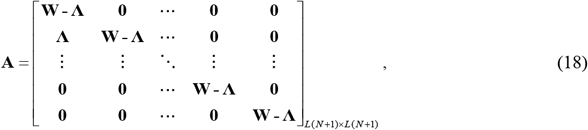

and the initial condition is

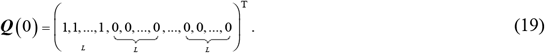

For *t* ≤ *t*, the solution of the above equation is

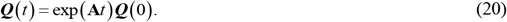

Therefore, we have

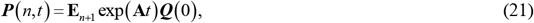

where

**E**_*n*_ = ***e***_*n*_ ⊗ ***e***, ⊗ represents the Kronecker production, ***e***_*n*_ denotes the *N*-dimension vector where the *n*-th coordinate is 1 and the others are zero, i.e., ***e***_*n*_ = (0,…,1,…,0)_1’(*N* +1)_, and ***e*** = (1,1,…,1)_1’*L*_ . For *t* > *t*_elo_, the solution of Eq. (3) is

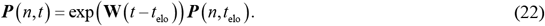

As *t* → ∞), we obtain the following matrix-form solution for the stationary nascent RNA distribution

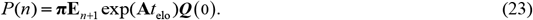

To verify the validity of the theoretical solution given by Eq. (23), we consider a four-state cyclic model of gene expression with deterministic elongation time. Fig. 1 (B)(C)(D) exhibits numerical [46] and theoretical results for the steady-state distribution of nascent RNA for three different sets of parameter values. From these diagrams, we observe that both numerical and theoretical results are well consistent.

### The two-state gene expression model: A case study

As a special case, we consider a general two-state model of gene expression with a deterministic elongation time, where we assume that the promoter switches between two states: an inactive state A and an active state I . The transcription initial rate is *k*_0_ at the I state and *k*_1_ at the A state. The two-state gene expression model is a special case of the cyclic model with *L*_1_ = 1 and *L*_2_ = 1 . The reaction scheme for this process is listed below

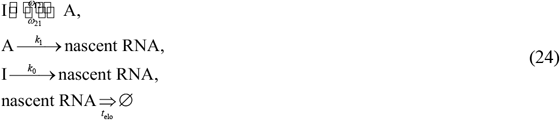

Previous studies have derived the analytical solution of nascent RNA distribution under the assumption that the transcription initiation only occurs at the active states [14,18], i.e., *k*_1_ > 0 and *k*_0_ = 0 . In this section, we derive the steady state distribution of nascent RNA for a general two-state gene expression model with leaky transcription, i.e., *k*_1_ > *k*_0_ ≥ 0 .

Applying Eq. (9), and using Sylvster’s matrix theorem to calculate [(*s***I** ‡ **U**)^‡1^ **Λ**]^*n*^, we can finally obtain the Laplace-form solution of nascent RNA distribution for the general two-state model as following

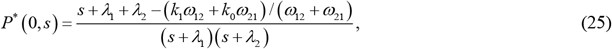

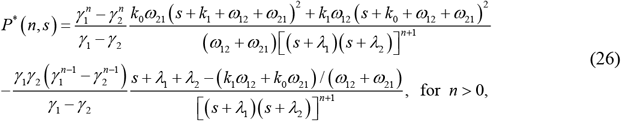

where ‡*λ*_1_ and ‡*λ*_2_ are roots of the quadratic equation

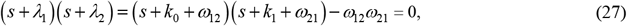

and *γ* _1_ and *γ* _2_ are roots of the quadratic equation

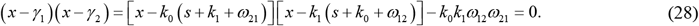

When *k*_0_ = 0, the roots of the above equation are *γ*_1_ = 0, *γ* _2_ = *k*_1_ (*s* + *ω*_12_) . Therefore,

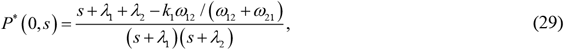

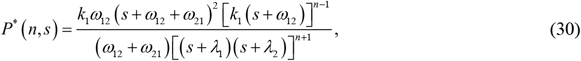

which reproduce the result in a previous study [18].

Using Eq.(13) and setting *τ* = *ω*_12_ + *ω*_21_, we obtain the Fanor factor of the nascent RNA number in the steady state

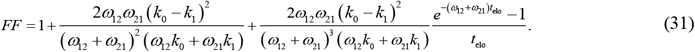

When *k*_1_ *>* 0 and *k*_0_ = 0, we have

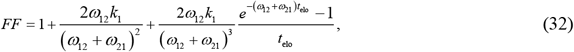

which reproduces the result in a previous study [23].

### Effect of molecular memory on nascent RNA distribution

Previous studies have demonstrated that molecular memory can modulate the distribution of mature RNA in both steady state and transient state [31-34]. However, the effect of molecular memory on the stationary distribution of nascent RNA remains incompletely understood. In this section, we investigate the influence of molecular memory on the stationary nascent RNA distribution using the above cyclic model. Note that this model considers two types of molecular memory: the memory during the active period (characterized by index *L*_1_ and called activation memory) and the memory during the inactive period (characterized by index *L*_2_ and called deactivation memory).

To understand how activation and deactivation memory affect the stationary nascent RNA distribution, we first focus on the cyclic model with fast or slow promoter switching. Under the slow-switching condition, i.e., *u* 0, *v* 0, it follows from Eq. (6) that *G* (*z*) = *π*_0_ exp(*k*_0_*t*_elo_ (*z* ‡1)) + *π*_1_ exp(*k*_1_*t*_elo_ (*z* ‡1)) with *π*_0_ = *v* / (*u* + *v*) and *π*_1_ = *u* / (*u* + *v*), indicating that the stationary nascent RNA distribution is a mixture Poisson distribution. In contrast, under the fast-switching condition, i.e., *u* →∞, *v* →∞, Eq. (6) becomes *G* (*z*) = exp((*π*_0_ *k*_0_ + *π*_1_*k*_1_)*t*_elo_ (*z* ‡1)), suggesting that the stationary nascent RNA distribution is a Poisson distribution. Therefore, both activation and deactivation memory do not alter the stationary distribution of nascent RNA in the limiting cases of fast and switching regimes.

We further investigate the impact of activation and deactivation memory on the stationary nascent RNA distribution in the case that promoter switching is neither too fast nor too slow. To illustrate this, we present the *u* ‡ *v* phase diagram for the two-state model with *L*_1_ = 1 and *L*_2_ = 1 (Fig. 2(A)), the multi-ON model with *L*_1_ = 5 and *L*_2_ = 1 (Fig. 2(B)) and the multi-OFF model with *L*_1_ = 1 and *L*_2_ = 5 (Fig. 2(C)). In all cases, the stationary nascent RNA distribution exhibits three main features: unimodal distribution with zero peak (Region I or Fig. 2(D)), unimodal distribution with non-zero peak (Region II or Fig. 2(E)), and bimodal distribution with a zero peak and a non-zero peak (Region III or Fig. 2(F)). Comparing these phase diagrams across the different models, we observe molecular memory can influence the size of each region: (1) increasing the activation memory *L*_1_ leads to the transition from unimodal distribution of type I to bimodal distribution, and results in a significant reduction of region I ; (2) increasing deactivation memory *L*_2_ causes the transition from bimodal distribution to the unimodal distribution of type II, and leads to an enlargement of Region II. In addition, we discover that increasing the deactivation memory *L*_2_ in the mult-OFF model induces a trimodal distribution with a zero peak and two non-zero peaks ((type IV or Fig. 2(G)), which has not been reported in previous studies.

**Fig. 2.**
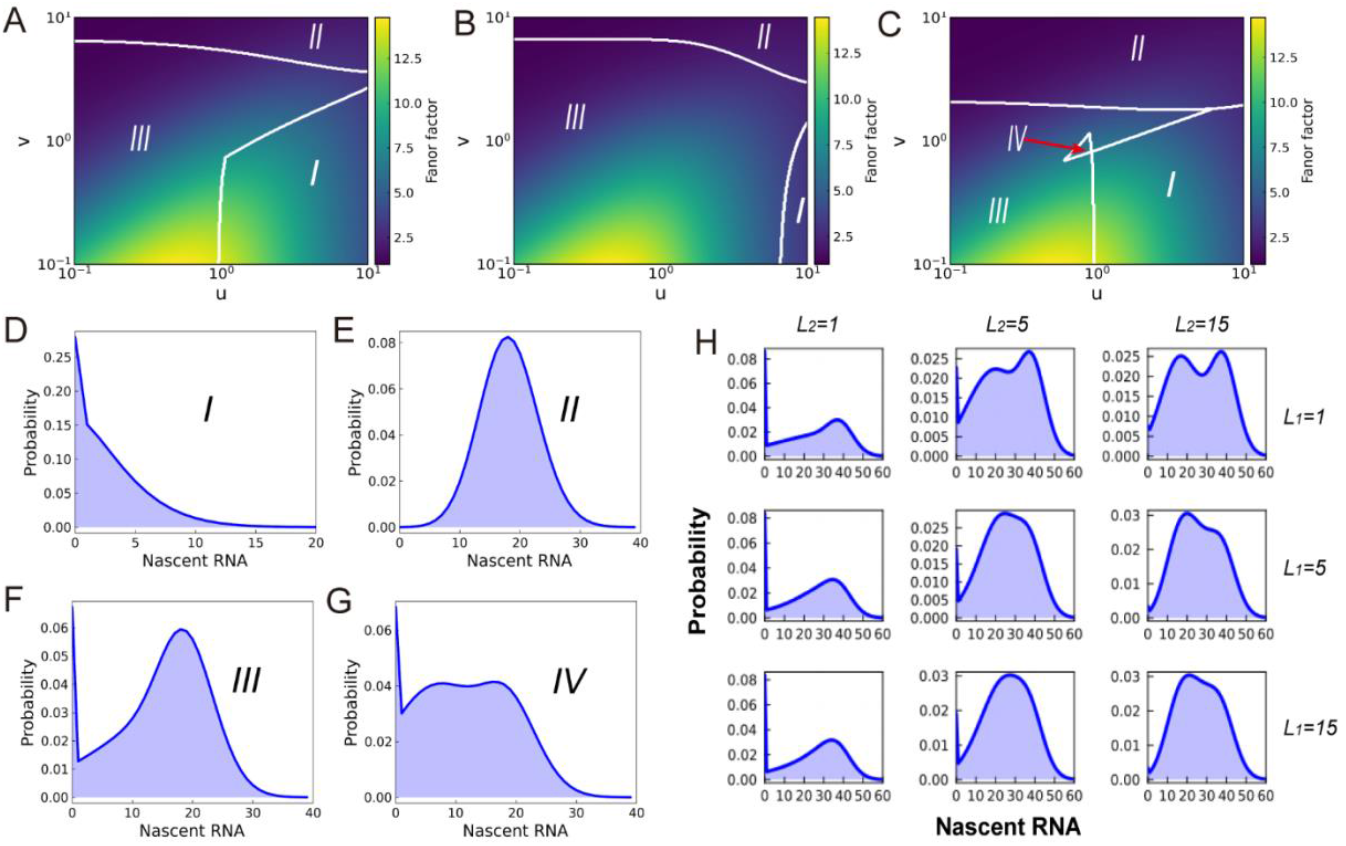
The effect of molecular memory on the nascent RNA distribution in the steady state. (A) The *u*-*v* phase diagram for the two-state model (*L*_1_ = 1, *L*_2_ = 1); (B) The *u*-*v* phase diagram for the multi-ON model (*L*_1_ = 5, *L*_2_ = 1) ; (C) The *u*-*v* phase diagram for the multi-OFF model (*L*_1_ = 1, *L*_2_ = 5); other parameter values are set as *k*_1_ = 20, *k*_0_ = 0; (D)(E)(F)(G) The distributions of different types, where the parameter values for each type are set as those for Type I *u* = 12, *v* = 2; Type II *u* = 1, *v* = 10; Type III *u* = 0.5, *v* = 1.5; Type IV *u* = 0.8, *v* = 1.2, *L*_2_ = 8. The other parameter values are the same as those in (A); (D) the steady state of nascent RNA distribution under different choices of *L*_1_ and *L*_2_. The other parameters values are set as: *u* = 0.8, *v* = 1.5, *k*_1_ = 40, *k*_0_ = 0.

In addition, we demonstrate stationary distributions of nascent RNA under different choices of *L*_1_ and *L*_2_, referring to Fig. 2(H). From this figure, we observe that increasing deactivation memory *L*_2_ tends to eliminate the zero peak, thus transforming the bimodal distribution into a unimodal distribution. When *L*_1_ = 1, increasing *L*_2_ can induce a new non-zero peak (the upper row in Fig. 2(H)), leading to the trimodal distribution in Region IV. When *L*_2_ is sufficient large, i.e., *L*_2_ = 15, the peak at zero almost disappears and the trimodal distribution becomes a bimodal distribution with two non-zero peaks. When both activation and deactivation memory are larger than 1, i.e., *L*_1_ > 1 and *L*_2_ > 1, two non-zero peaks of the bimodal distribution tend to merge together, leading to a unimodal distribution.

To understand why increasing the activation memory *L*_1_ can transform the unimodal distribution of type I to the bimodal distribution, we investigate the stationary nascent RNA distribution in the burst regime (*u* ▯ *v*). From Fig. 3(C)(D), we observe that increasing activation memory *L*_1_ diminishes the probability *P* (*n* = 0) and *P* (*n* = 1) simultaneously, concurrently giving rise to a valley between two peaks. This implies that activation memory *L*_1_ can induce bimodal distribution in the burst regime. However, within the burst regime, increasing *L*_2_ significantly reduces the probability *P* (*n* = 0) while increase the probability *P* (*n* = 1) (Fig. 3(A)(B)), and the unimodal distribution remains unchanged. Hence, the activation and deactivation memories exhibit distinct modulation effects on nascent RNA distribution. It should be noted that this characteristic may serve as an indicator for distinguishing activation memory form station nascent RNA distribution, i.e., when the stationary nascent RNA displays bimodal distribution in the bursting regime, it suggests that the activation memory may be larger than 1.

**Fig. 3.**
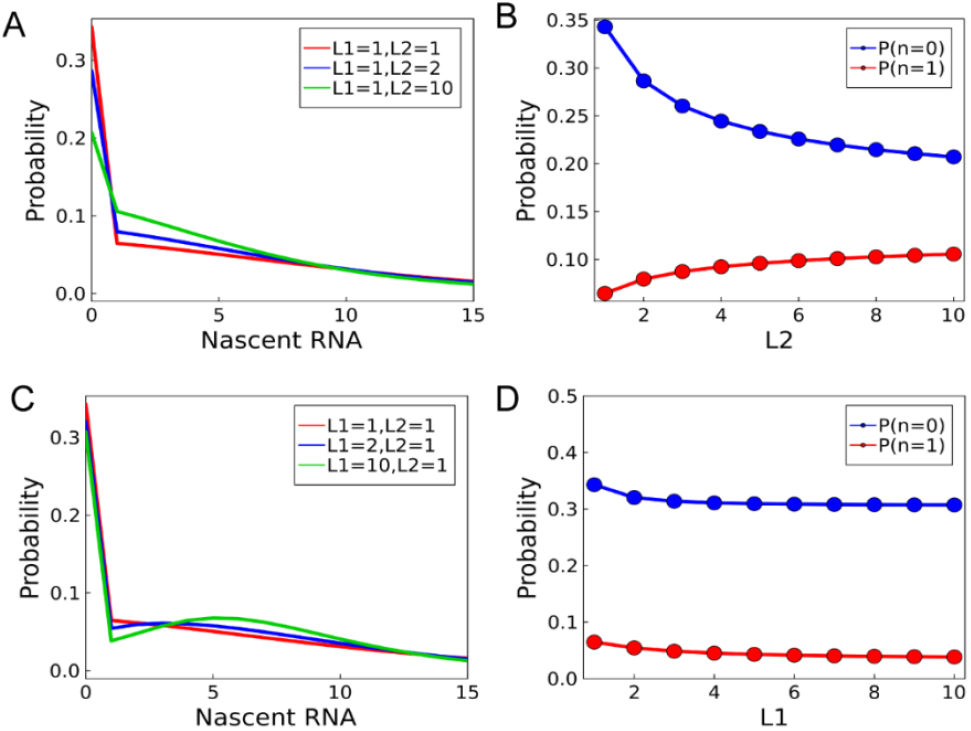
The effect of molecular memory on the stationary nascent RNA distribution in the burst regime (*u* ▯ *v*) . (A) The steady-state distributions of nascent RNA for different values of *L*_2_ ; (B) The probabilities of *P* (*n* = 0) and *P* (*n* = 1) as functions of *L*_2_; (C) The steady-state distribution of nascent RNA for different value of *L*_1_; (D) The probabilities of *P* (*n* = 0) and *P* (*n* = 1) as functions of *L*_1_ . The other parameter values are set as: *u* = 3, *v* = 1, *k*_1_ = 20, *k*_0_ = 0.

By summarizing the above analysis, we know that the molecular memories during active and inactive periods can modulate the steady-state distribution of nascent RNA cooperatively. Increasing the deactivation memory *L*_1_ tends to disrupt the zero peak in the stationary nascent RNA distribution, while enhancing the activation memory *L*_2_ can induce a bimodal distribution in the bursting regime. Depending on the sizes of the activation and deactivation memories, distinct shapes of stationary nascent RNA distribution may emerge, including bimodal distribution with a zero peak and non-zero peak, bimodal distribution with two non-zero peaks, and trimodal distribution, unimodal distribution with zero peak, and unimodal distribution with non-zero peak.

### Inferring the memory of promoter switching from a nascent RNA distribution

The nascent RNA distribution can be obtained through experimental techniques such as mRNA-FISH labeling or single-cell RNA sequencing (scRNA-seq). An intriguing question is whether the memory of promoter switching (called promoter memory for convenience) can be inferred from the stationary nascent RNA distribution. The question of how model parameters are inferred from a given distribution has been explored in previous studies [47-50]; and the two-state non-Markov model has commonly been employed to fit the nascent RNA distribution and to elucidate the underlying mechanism of gene expression [14,15]. In this section, we propose a method based on the above analytical results to infer the promoter memory from the stationary nascent RNA distribution.

We assume that *M* cells are measured simultaneously at an one-time experiment and cell *i* has *n*_*i*_ copies of nascent RNA in the steady state, and the stationary distribution of nascent RNA is *P*(*n,θ*) . We also assume that the promoter consists of *L*_1_ active states and *L*_2_ inactive states. The transition rates at the ON (OFF) states are the same, i.e., 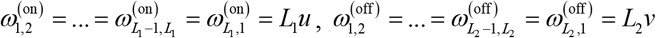. The initiation of nascent RNA only occurs in the active state with rate *k*_1_ . Since it is impossible to infer the elongation time and other parameters simultaneously from a stationary distribution, we fix the elongation time *t*_elo_ = 1 . Let *θ* = (*L*_1_, *L*_2_,*u, v, k*_1_) be the parameter vector of the cyclic model to be inferred. We define the total likelihood function as

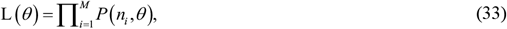

where *P* (*n* ;*θ*) is calculated according to Eq. (23). We use an efficient gradient-free optimization algorithm (i.e., adaptive differential optimizer, referring to BlackBoxOptim.jl [51]) to find the optimal parameters by maximizing the likelihood function L (*θ*) .

To evaluate the accuracy of the method, we generate a set of synthetic data comprising 2000 cells using the stochastic simulation method [46]. It is worth noting that the parameter set for this synthetic data is sampled in the burst regime, satisfying the condition *u* ▯ *v* . By fitting both the cyclic model and classical two-state model to this synthetic data, we observe that the former outperforms the latter in terms of goodness of fit (Fig. 4(A)). In order to assess estimation accuracy, bootstrapping resampling (i.e., random sampling with replacement) is performed to obtain a distribution of the estimated parameters, which can be utilized to estimate the standard errors and the confidence intervals of these estimated parameters [52]. As shown in Fig. 4(B), the activation memory *L*_1_ can be accurately estimated. Furthermore, Fig.4(C) demonstrates that all ‘true’ parameter values fall within the 95% confidence interval estimated by the bootstrapping, indicating the precise inference of the ‘true’ parameters from the stationary nascent RNA distribution in the burst regime. However, extensive numerical simulations reveal limitations in accurately inferring promoter memory solely based on stationary nascent RNA distribution under certain conditions. For instance, when comparing promoter switching rates with elongation process rates—either sufficiently fast or slow— the nascent RNA distribution becomes independent of promoter memory.

**Fig. 4.**
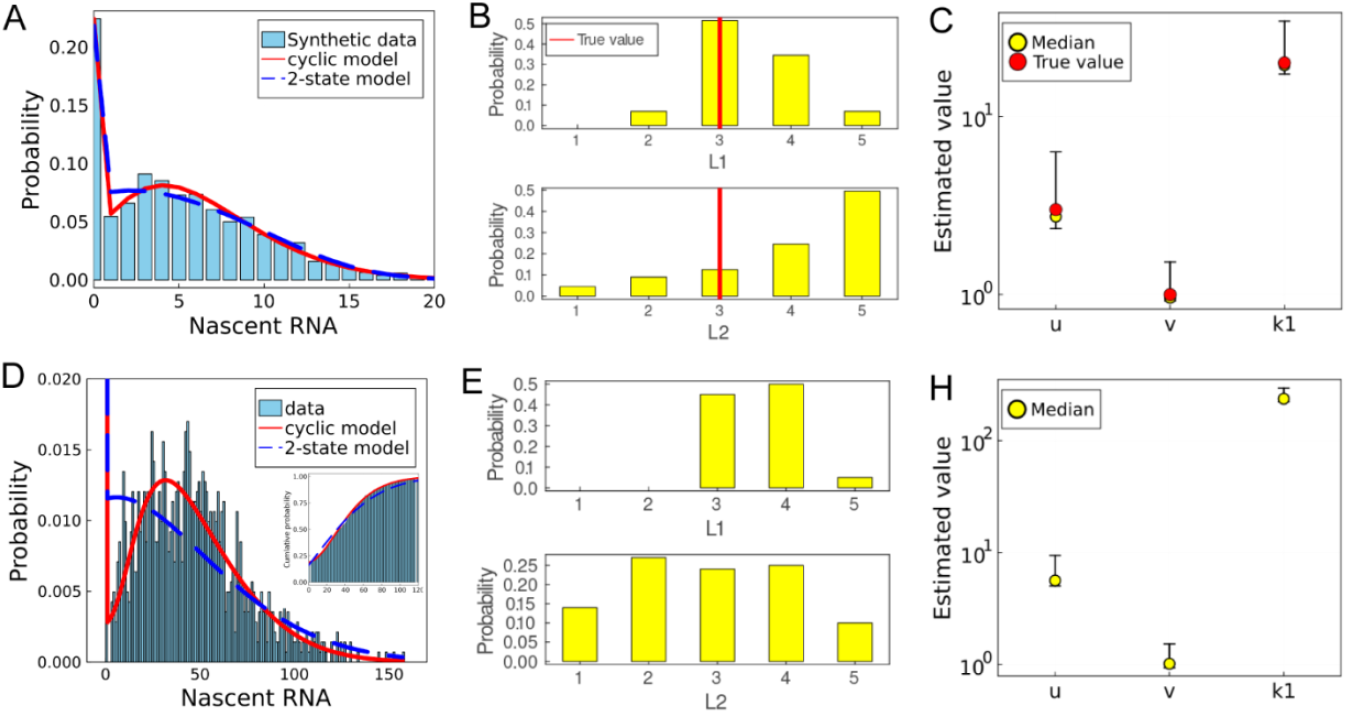
Inference of promoter memory from nascent RNA distribution. (A)The fitted distributions for synthetic data using the cyclic model and the two-state model. The ‘true’ parameter set for the synthetic data is: *L*_1_ = 3, *L*_2_ = 3, *u* = 3, *v* = 1, *k*_1_ = 20 ; (B) the distribution of promoter memory estimated by bootstrap resampling for the synthetic data. The red line represents the ‘true’ value; (C) the 95% confidence interval of the kinetic parameters estimated by bootstrap resampling for the synthetic data. The red circle represents the ‘true’ value. (D) The fitted distributions for experimental data of *HIV-1* gene using the cyclic model and the two-state model. The estimated parameters of the cyclic model are: *L*_1_ = 3, *L*_2_ = 4, *k*_1_ = 242.66, *u* = 5.12, *v* = 0.95 ; the estimated parameters of the two-state model are: *k*_1_ = 122.21, *u* = 3.41, *v* = 1.50 . The inset is the cumulative probability function that corresponds to the distribution estimated by the cyclic and two-state models; (E) the distribution of the promoter memory estimated by bootstrap resampling for the experimental data; (H) the 95% confidence interval of kinetic parameters estimated by bootstrap resampling for the experimental data.

Finally, we apply the aforementioned inference method to a realistic example. Specifically, we aim to infer the promoter memory from the stationary nascent RNA distribution for the *HIV-1* gene in cells expressing a high level of Tat which is measured by Katjana et.al. [20]. It has been reported that the expression of the *HIV-1* gene exhibits bursting kinetics, which are believed to play a crucial role in the control of latency. We fit the experimental data using the two-state model and cyclic model, and present the distribution of the parameters estimated by bootstrapping resampling in Fig. 4(E)(H). Notably, this analysis reveals that the activation memory *L*_1_ is larger than 1. By comparing with Fig. 4(D), we find that our proposed cyclic model outperforms the two-state counterpart in terms of fitting accuracy. Furthermore, although inferring an exact distribution of the dwell time at the active state remains challenging, our estimations strongly suggest non-exponential behavior of the dwell time during activation.

## Conclusions and discussion

In this study, we have analyzed a general stochastic model of gene expression with complex promoter structure, which included many previously studied models as its special case, and derived analytical results for the stationary distribution and statistical quantities of nascent RNA, which provide insight into the stochastic kinetics of nascent RNA. Interestingly, we found that the molecular memories created by multistep activation and deactivation can modulate the steady-state distribution of nascent RNA cooperatively, e.g., increasing the deactivation memory tends to disrupt the zero peak in the stationary nascent RNA distribution, while enhancing the activation memory can induce a bimodal distribution in the bursting regime. More practically, we developed an efficient method to infer the shape parameters characterizing the non-exponential distributions of the waiting times between promoter states from stationary nascent RNA distributions, and verified the validity of our inference method by analyzing the experimental data of a realistic example.

Using the analytical solution of a cyclic model, we investigated how activation and deactivation memory affect the shape of the stationary nascent RNA distribution. Previous study [14] showed that the two-state model with deterministic elongation time can display bimodal distribution with a zero peak and non-zero peak. By contrast, our study revealed that increasing the deactivation memory can destroy the zero peak, resulting in a bimodal distribution with two non-zero peaks. Additionally, we found that increasing the initiation rate *k*_0_ at the inactive state also eliminated the zero peak. This suggests that the zero peak in stationary nascent RNA distribution is unstable, and this fact may explain why the bimodal distribution is rarely observed in experiments. Furthermore, our study showed that by increasing activation memory, delay-induced bimodality or delay-induced zero-inflation phenomenon for nascent RNA can be further enlarged within burst switching regime as reported by Qingchao et al. [17]. However, our study has limitations since it only considers a cyclic promoter proceeding sequentially through several irreversible active and inactive state while real-life promoters may have more complex structure such as competitive cross-talking pathways [53] and reversible processes [31], all of which may affect stationary nascent RNA distributions.

Based on the analytical solution of the stationary nascent RNA, we have also developed an effective method to infer the promoter memory and transcriptional burst kinetics from a known nascent distribution. Our inferred results indicated that in the burst regime where the promoter spends most of its time in the inactive period, it is possible to infer promoter memory, in particular active memory, from the stationary nascent RNA distribution. We applied this method to the *HIV-1* gene [20], which was reported to exhibit bursting kinetics. By fitting the experimental data to the cyclic model, we found that the active memory was always larger than 1, suggesting a non-exponential dwell time distribution during activation. However, it should be noted that accurately inferring promoter memory from stationary nascent RNA distribution may not always be feasible since different promoters with distinct active and inactive memories can generate identical stationary distribution [31,54]. Further investigation is required to address potential bias in inferring promoter memory from nascent RNA distribution. Additionally, intermediate processes of gene expression such as the partitioning of mRNA/protein during cell division [5,55,56], would contribute the noise of gene expression and potentially skew the inferred results, but these processes were not considered in our study. How details of intermediate processes affect the inferred results is worth further study.

## Acknowledgments

This work was supported by grants 12001129, 12171494, 62373384, 11931019, and 12371483 from the Natural Science Foundation of P. R. China, and by grants 2019B110233002, 202007030004 from Key-Area Research and Development Program of Guangzhou, P.R. China, by grants 2022ZDZX2045 from the Special Projects in Key Fields for Colleges and Universities in Guangdong Province.

## Data Availability

The nascent RNA data of *HIV-1* gene in cells expressing a high level of Tat were downloaded from ref. [20].

